# FAM19A4 Enhances Neutrophil Respiratory Burst via p38 MAPK in Lethal Sepsis

**DOI:** 10.1101/2025.11.07.687190

**Authors:** Shu Li, Fengxue Zhu, Lilei Jiang, Haiyan Xue, Ting Li, Tianbing Wang, Wenyan Wang, Kai Zhang

**Author notes:** **Correspondence:** Fengxue Zhu.

## Abstract

Sepsis causes high mortality and resource strain, with neutrophil-derived reactive oxygen species (ROS) contributing to excessive inflammation. The secretory protein FAM19A4 modulates ROS release, but its role in sepsis was unclear. We measured elevated FAM19A4 levels in septic patients and cecal ligation and puncture (CLP) mice, correlating with increased mortality. *Fam19a4^−/−^* mice subjected to CLP showed significantly improved survival and attenuated multiorgan injury without impaired peritoneal bacterial clearance or altered circulating neutrophil counts. Deficiency reduced the cell counts of neutrophil (Ly6G^+^) and macrophage (F4/80^+^) in lungs and liver, diminished systemic ROS production tracked by bioluminescence, and decreased neutrophil extracellular trap (NET) formation in serum and lung tissue. In vitro, FAM19A4 enhanced neutrophil phagocytosis and ROS generation but did not affect lipopolysaccharide-induced chemotaxis. Mechanistically, bulk RNA sequencing, western blotting, and the p38 inhibitor SB203580 demonstrated that FAM19A4 drives neutrophil ROS release specifically through p38 MAPK signaling activation. These results indicate that FAM19A4 is upregulated during sepsis and exacerbates outcomes by enhancing neutrophil ROS production via p38 MAPK, representing a promising therapeutic target for this condition.

## 1 Introduction

Sepsis is a life-threatening organ dysfunction caused by a dysregulated host response to infection [1]. With an estimated incidence of 18.6 cases per 1000 hospitalizations and mortality rates approaching 40% [2], sepsis imposes substantial long-term healthcare burdens on survivors while straining global medical resources [3, 4]. The pathophysiology of sepsis involves hyperactivation of innate immunity, characterized by excessive cytokine production (“cytokine storms”), that drives cardiovascular collapse, multiorgan failure, and early mortality [5]. Despite progress, key molecular drivers of this dysregulated response remain undefined, hampering development of targeted diagnostics and therapies. Elucidating the mechanisms underlying sepsis development represents a critical frontier for advancing management of this condition.

Neutrophils are crucial mediators of dysregulated inflammation during the early innate immune phase of sepsis [6]. Especially in sepsis-associated acute respiratory distress syndrome (ARDS), neutrophils, as first responders, are among the earliest and most abundant immune cells recruited to the lung [7]. While neutrophils are essential for microbial clearance, their pathological accumulation in vital organs triggers respiratory burst-derived reactive oxygen species (ROS) overproduction [8], and these oxygen radicals induce mitochondrial dysfunction and propagate cellular damage cascades [9]. Notably, activated neutrophils release DNA scaffolds decorated with histones, granule proteins, and antimicrobial peptides, referred to as neutrophil extracellular traps (NETs) [10]. ROS generation is a critical driver of NET formation via NADPH oxidase and myeloperoxidase (MPO) [11]. Excessive NETosis exacerbates tissue injury through proinflammatory cascades, and is correlated with sepsis-associated mortality rates [12, 13]. p38 mitogen-activated protein kinase (p38 MAPK), a core member of the stress-activated MAP kinase superfamily, undergoes phosphorylation-mediated activation at a conserved TGY motif within its kinase activation loop, and this process is initiated by proinflammatory cytokines and mitogens through sequential kinase cascades [14]. p38 MAPK potentiates neutrophil ROS generation by enhancing NADPH oxidase activation in response to proinflammatory stimuli and pathogen-associated molecular patterns; critically, this pathway directly mediates sepsis-induced organ injury through ROS-induced tissue damage [15].

FAM19A4 (family with sequence similarity 19, member A4), also referred to as TAFA4, is a member of the TAFA protein family [16]. Human FAM19A4 is widely expressed at low levels in various normal tissues and organs, while in mouse, FAM19A4 is only highly expressed in the dorsal ganglia [17]. Lipopolysaccharide (LPS)-stimulated FAM19A4 potentiates phagocytic capacity and ROS generation [18]. Further, FAM19A4 directly modulates the in vitro inflammatory profile of macrophages from a mouse sunburn-like model of skin damage and promotes IL-10 production by dermal macrophages in vivo [19]. Given its immunomodulatory properties and association with myeloid cell function, FAM19A4 could potentially be a critical regulator of sepsis pathophysiology; its impact on neutrophils (central effectors of innate immunity in sepsis) warrants particular investigation. In this study, we examined the organ-protective and prognostic benefits of FAM19A4 deficiency in sepsis and elucidated mechanistic insights in the context of ROS release.

## 2 Materials and Methods

### 2.1 Generation of *Fam19a4*-EGFP knock-in (KI) mice

To generate *Fam19a4*-EGFP KI mice, the EGFP-WPRE-pA expression cassette was inserted through CRISPR/Cas9-mediated homologous recombination, precisely 30 bases downstream of the *Fam19a4* gene start codon. *Cas9* mRNA and single-guide RNA (gRNA) were produced by in vitro transcription. A donor vector for homologous recombination was constructed containing a 4.1-kb 5’ homology arm, the EGFP-WPRE-pA cassette, and a 2.8-kb 3’ homology arm using In-Fusion cloning. Subsequently, *Cas9* mRNA, gRNA, and the donor vector were co-injected into fertilized C57BL/6 oocytes to produce F0 founder mice. Resulting chimeras were bred with wild-type (WT) C57BL/6 females to achieve germline transmission of the targeted allele. Genotyping was performed using quantitative PCR (qPCR). For all experiments, age- and sex-matched littermates derived from the same breeding pairs were utilized, and all animals were maintained on a pure C57BL/6 genetic background.

### 2.2 Cecal ligation and puncture (CLP)-induced polymicrobial sepsis model

Sepsis was induced in mice (20-22 g body weight) using a previously described CLP model [20]. Following skin disinfection with 0.3% chlorhexidine, a midline laparotomy was performed under 2% isoflurane anesthesia delivered in oxygen. The cecum was ligated at its distal 25% segment and was subsequently punctured once with a 20-gauge needle. Postoperative resuscitation included subcutaneous administration of 1 mL sterile saline. Humane endpoints were enforced through CO_2_ euthanasia for moribund animals or at 7 days (experiment termination). In the control group, animals underwent anesthesia and laparotomy without cecal ligation or puncture.

### 2.3 Isolation of human neutrophils

Human neutrophils were isolated from healthy adult volunteer venous blood samples, as described previously [21]. Venous blood collected in EDTA-coated vacutainers was diluted with 20 mL PBS and under layered with lymphocyte separation medium (8 mL), followed by centrifugation (1500 rpm, 25 min; brake off). The RBC/neutrophil pellet was retained, resuspended in 20 mL Hank’s balanced salt solution and 20 mL 3% dextran/0.9% NaCl, then incubated for 30 min at 20°C. The neutrophil-rich upper layer was transferred and centrifuged (400×g, 8 min), followed by sequential osmotic lysis of the pellet by resuspension in 0.2% NaCl, followed by 1.6% NaCl addition, with centrifugation (400 × g, 8 min) after each step. Finally, pellets were resuspended in RPMI + 14% fetal bovine serum (FBS). Neutrophil counts, viability (trypan blue exclusion), and purity (> 99% via Giemsa staining) were confirmed.

### 2.4 FAM19A4, IL-6, IL-10 and TNF-**α** measurement

Peripheral blood samples from mice and patients with sepsis were centrifuged (3000 rpm, 5 min) to collect serum for analysis. Bronchoalveolar lavage fluid (BALF) was obtained by clinicians following standardized protocols, as follows: under local anesthesia, 100 mL pre-warmed (37°C) sterile saline was instilled into the right middle lobe or left lingula in 30-50 mL aliquots via electronic fiberoptic bronchoscopy; lavage fluid was aspirated under 50-100 mmHg negative pressure (minimum 40% recovery rate), centrifuged (800 rpm, 10 min), and the supernatant retained. Cytokine levels (IL-6, IL-10, TNF-α) in serum and BALF were quantified using eBioscience ELISA kits, while FAM19A4 was measured with MyBioSource kits, per manufacturers’ protocols.

### 2.5 In situ hybridization

In situ hybridization was carried out with a digoxigenin-labeled probe, which was hybridized for 20 hours at 50°C. We then probed the liver and lung sections with a horseradish peroxidase (HRP)-labeled anti-digoxigenin antibody (Roche, Shanghai, China). Signal visualization was accomplished using a fluorescein/cy3/cy5 TSA plus amplification kit (Perkin Elmer Inc., Beijing, China).

The following oligonucleotides were used for probe synthesis by nested PCR:

*Fam19a4*-F1: TGCTCAGAAGTTCATAGCCAAA

*Fam19a4*-R1: TAAAGGAACATTTGCAAGCTCA

*Fam19a4*-F2: ATATGTGCAGTGTGG

*Fam19a4*-R2+T7: TAATACGACTCACTATAGGGCAGCCAAGTTCAAAC

### 2.6 Histopathology

For subsequent histopathological assessment, tissues were subjected to fixation in 10% neutral-buffered formalin and subsequent processing through standardized protocols for paraffin embedding and sectioning. Serial 2-μm sections were stained with hematoxylin & eosin (H&E), anti-F4/80 (AbD Serotec; clone CI:A3-1), or anti-Ly6G (BioLegend; clone 1A8). H&E-stained sections underwent microscopic evaluation with semi-quantitative scoring, as follows: renal tubular necrosis (0: no injury; 1: 0%–25%; 2: 25%–50%; 3: 50%–75%; 4: > 75% affected tubules) [22], lung injury assessed via Nishina score [23], and hepatic pathology graded using Suzuki criteria [24]. For immunohistochemical staining, primary antibodies were used at 1:100 dilution following antigen retrieval (BOND Epitope Retrieval kit, Leica, Beijing, China) with DAB visualization. The quantity of Ly6G^+^ and F4/80^+^ cells in kidney, liver, and lung tissues was evaluated using Definiens Tissue Studio software (Definiens, Inc.). Representative regions were manually annotated to train artificial intelligence algorithms for target object identification (morphology/positivity-based) and background exclusion. Validated algorithms were subsequently applied for full-slide automated analysis, with diagnostic concordance confirmed by blinded histopathological review.

### 2.7 Quantitative PCR

Total RNA was isolated from liver, lung, kidney, and spleen tissues using RLT Plus buffer with β-mercaptoethanol homogenization, followed by centrifugation (12,000 × g, 3 min, 4°C). After genomic DNA removal via DNase I digestion on Omega Bio-tek columns, RNA was purified through RW1 buffer washes and eluted in RNase-free water, achieving spectrophotometric purity (A□□□/A□□□=1.8–2.1; A□□□/A□□□>2.0). Reverse transcription of 1 µg RNA using PrimeScript RT Kit with random hexamer/oligo(dT) priming generated cDNA, which underwent quantitative PCR with PowerUp™ SYBR™ Green Master Mix on QuantStudio 5 under optimized cycling: UDG activation (50°C, 2 min), denaturation (95°C, 2 min), 40 cycles of 95°C/15s and 60°C/1min, followed by melt curve analysis. Relative gene expression was calculated using the 2^-ΔΔCT^ method normalized to *Gapdh*. The following primer sequences were used for amplification:

*Fam19a4* forward: ATGAGGTCCCCAAGGATGAGAGTCTG

*Fam19a4* reverse: CTCACAGGTCCCTTGCTTGATTTGGT

*Gapdh* forward: GTGGCAAAGTGGAGATTGTTG

*Gapdh* reverse: CGTTGAATTTGCCGTGAGTG

### 2.8 Phagocytosis assay

Neutrophil phagocytic function was assessed using a pHrodo *E. coli* Bioparticles Phagocytosis Kit (Life Technologies, CA, USA). Neutrophils were pre-treated with varying concentrations of FAM19A4 protein, either alone or in combination with LPS, followed by incubation with pHrodo *E. coli* bioparticles for 2 hours at 37□°C. Phagocytosis was quantified by either confocal microscopy (Leica TCS SP8, Wetzlar, Germany) or flow cytometry using a FACSVerse system (BD Biosciences, San Jose, CA, USA) and BD FACSuite software. For flow cytometric analysis, the percentage of Ly6G^+^ cells exhibiting pHrodo *E. coli* positivity was determined based on a gating strategy established from unstimulated versus bioparticle-treated cells. Fluorescence emission occurred following the internalization of particles into acidic phagosomal compartments. For microscopy-based quantification, three representative fields were imaged, and the average number of internalized pHrodo *E. coli* particles per neutrophil was calculated from ten randomly selected Ly6G^+^ cells.

### 2.9 Western blotting

Neutrophils were rinsed twice with ice-cold PBS. Cell lysis was carried out using RIPA buffer containing protease and phosphatase inhibitor cocktails. The resulting lysates were combined with 3×SDS loading buffer, subjected to heat denaturation, and then loaded onto 8% SDS-PAGE gels. Proteins (25 μg per lane) were separated electrophoretically on 8% polyacrylamide gels with Tris-glycine running buffer and subsequently transferred onto nitrocellulose membranes. The membranes were incubated sequentially with specific primary antibodies and HRP-conjugated secondary antibodies at a dilution of 1:5,000. Detection was performed with enhanced chemiluminescence (ECL) substrate using a FluorChem FC3 imaging system (ProteinSimple, USA), and band intensities were quantified with AlphaView software (version 3.4.0). Levels of phosphorylated p38 were normalized to total p38 protein and expressed as a percentage of the control values.

### 2.10 Bioluminescence imaging

Mice were kept anesthetized through sustained isoflurane inhalation and oriented dorsally for image acquisition. Image capture was conducted on an IVIS 200 platform (Xenogen Corporation, Alameda, CA), which was set up for luminescence detection without an optical filter and with the aperture fixed at F/Stop 1. For every imaging session, L-012 was delivered intravenously at 25 μg per gram body weight, using a solution of 20 μg/μl in sterile PBS. Quantification of signals from user-defined regions (photon flux) was executed within Living Image software (v.4.2, Xenogen Corporation) [25]. Statistical comparisons across time points were performed via two-way ANOVA supplemented with Bonferroni’s post hoc analysis.

### 2.11 Quantification and visualization of NETs

Levels of circulating NET components, such as citrullinated histone 3 (CitH3), MPO, and DNA, were quantified using established CitH3-DNA and MPO-DNA complex ELISA assays according to a previously reported method [26]. Briefly, 96-well plates pre-coated with capture antibodies (anti-CitH3 or anti-MPO, Abcam, Beijing, China) were blocked, then incubated with serum samples and subsequently with peroxidase-conjugated anti-DNA detection antibody. After tetramethylbenzidine substrate development, absorbance was measured at 450 nm. NET levels are expressed as fold change relative to control samples.

For NET visualization, lung tissue sections were subjected to immunostaining using specific primary antibodies incubated overnight at 4°C. Afterward, sections were treated with FITC-conjugated secondary antibodies (Abcam, Beijing, China) for 1 hour at room temperature. Nuclei were labeled with DAPI-containing mounting medium, and fluorescent images were obtained using a microscope. Both DAPI and FITC signals from the same tissue regions were collected to allow colocalization analysis.

### 2.12 Determination of superoxide release

Extracellular ROS generation by neutrophils was assessed via the reduction of ferricytochrome c. Neutrophils (10^6^ cells/ml) were incubated with 0.5 mg/ml ferricytochrome c and 1 mM Ca^2+^ at 37°C, pretreated with LPS (10 μg/ml) for 3 minutes and cytochalasin B (1 μg/ml) for 1 minutes, and then stimulated with varying concentrations of FAM19A4 (1, 10, and 100 ng/ml) for 5 minutes. Absorbance changes were monitored continuously at 550 nm using a U-3010 spectrophotometer (Hitachi, Tokyo, Japan).

### 2.13 Total ROS release measurement

ROS were measured using luminol-enhanced chemiluminescence. Neutrophil suspensions (1×10^6^ cells/mL) were first incubated with 35 μM luminol and 5 U/mL HRP for 5 minutes at 37°C, then stimulated with LPS (10 μg/mL) for another 3 minutes. Subsequently, cells were exposed to FAM19A4 at concentrations of 1, 10, or 100 ng/mL for 8 minutes. Real-time chemiluminescence signals were recorded on a Tecan Infinite F200 Pro multimode microplate reader (Tecan Group, Männedorf, Switzerland).

### 2.14 Intracellular ROS Production Assay

Intracellular ROS levels were quantified with a commercial assay kit utilizing the fluorescent indicator DCFH-DA. Peripheral blood mononuclear cells (PBMCs) were obtained from CLP-modeled mice by density gradient centrifugation with Ficoll-Paque PLUS (GE Healthcare). In brief, blood samples were diluted 1:1 with PBS, carefully layered onto the density gradient medium, and centrifuged at 400×g for 20 minutes at room temperature. The PBMC band was harvested, washed twice with PBS, and resuspended in RPMI-1640 medium at a density of 1×10^6^ cells/mL. Subsequently, cells were loaded with 10μM DCFH-DA and incubated for 30 minutes at 37 °C. After incubation, unincorporated dye was removed by washing, and cells were stimulated with cytochalasin B (CB; 0.5 μg/mL) for 2 minutes to potentiate ROS generation. The reaction was stopped by adding ice-cold PBS supplemented with 2% FBS and 1 mM EDTA. Samples were promptly analyzed on a BD FACSCanto II flow cytometer using a 488 nm excitation laser and a 530/30 nm emission filter. A total of 10,000 viable cells, gated based on forward and side scatter properties, were acquired per sample. Mean fluorescence intensity (MFI) of DCF-positive cells was determined using FlowJo software.

### 2.15 Bulk RNA sequencing

Total RNA was isolated from human neutrophils with Trizol reagent. RNA purity was assessed using a NanoPhotometer spectrophotometer (IMPLEN, Munich, Germany), and integrity was evaluated with a bioanalyzer (Agilent). The DNBSEQ platform was employed for RNA sequencing and subsequent bioinformatic processing.

Differentially expressed genes (DEGs) were screened for based on relative expression and fold change thresholds (log2(Fold Change) ≥ 0.5 and P < 0.05) and selected for further analysis. Volcano plots, heatmaps, and KEGG pathway enrichment were constructed using the online tool suite at http://www.omicstudio.cn/tools. Volcano plots were plotted from statistically significant DEGs. Intersecting gene sets were identified, and candidate sequences were chosen according to KEGG enrichment results. Heatmaps were generated to display expression patterns of DEGs.

### 2.16 Ethics approval and consent to participate

Animal experiments were conducted in accordance with the principles outlined in the Animal Experimentation Ethics Committee Guide for the Care and Use of Laboratory Animals and were approved by the Animal Experimentation Ethics Committee of Peking University Peoples’ Hospital (No. 2019PHE030). Experimental protocols involving human data and tissue were approved by the Institution of Human Subjects Committee at Peking University People’s Hospital (No. 2019PHB191-01). Informed written consent was obtained from all subjects.

### 2.17 Data and statistical analysis

Figures and statistical analyses were prepared and performed with GraphPad Prism software. Histopathological evaluations were carried out under blinded conditions using Definiens Tissue Studio. Mice were assigned to experimental groups according to genotype and randomized within cohorts matched by sex and age. Group sizes were determined by empirical assessment of variability within each model or assay system, with the number of mice per group maximized to reduce both type I and type II errors.

Data with a normal distribution are expressed as mean ± SD, and non-normally distributed data as median ± interquartile range (IQR). Since all datasets in this study passed normality testing, results are reported as mean ± SD to reflect experimental variance. Significance of differences was assessed using the following tests: the non-parametric log-rank test for survival data; for parametric data, the two-tailed unpaired Student’s t-test was used for comparisons between two groups, and one-way ANOVA followed by Tukey’s multiple comparisons test was used for comparisons among more than two groups. All parametric data were confirmed to be normally distributed by the Shapiro-Wilk test as stated above. Significance notations are: “ns” for not significant, *P < 0.05, **P < 0.01, and ***P < 0.001. For gene enrichment analysis, statistical significance was evaluated using the Q-value, which represents the false discovery rate-adjusted p-value.

## 3 Results

### 3.1 FAM19A4 expression is elevated in patients with sepsis and CLP-subjected mice

Under physiological conditions, FAM19A4 expression in peripheral tissues (liver, lung, kidney, intestine) is minimal, with higher levels detected in the central nervous system [16]. GEO Profiles data (GDS2856/GPL570 probe 242348_at; GDS2430/GPL97 probe 242348_at) indicate that LPS induces upregulation of FAM19A4 in monocytes/macrophages. In vitro, LPS stimulation promotes FAM19A4 secretion from M1-polarized macrophages and monocytes [18]. To characterize FAM19A4 expression during sepsis hyperinflammation, we enrolled 40 patients meeting Sepsis-3.0 criteria (≤ 7 days post-onset) and 10 healthy volunteers as the control group. Serum and BALF from patients and volunteers were analyzed via ELISA, revealing significant FAM19A4 elevation in patients with sepsis vs. healthy volunteers (Figure 1A). Analysis of 28-day mortality revealed lower FAM19A4 concentrations in survivors than those in non-survivors (Figure 1B). FAM19A4 levels correlated with both inflammatory response intensity and mortality risk, while they were almost undetectable in healthy volunteers. ELISA quantification of IL-6, TNF-α, and IL-10 via served as reference biomarkers for systemic inflammation [25].

**Figure 1.**
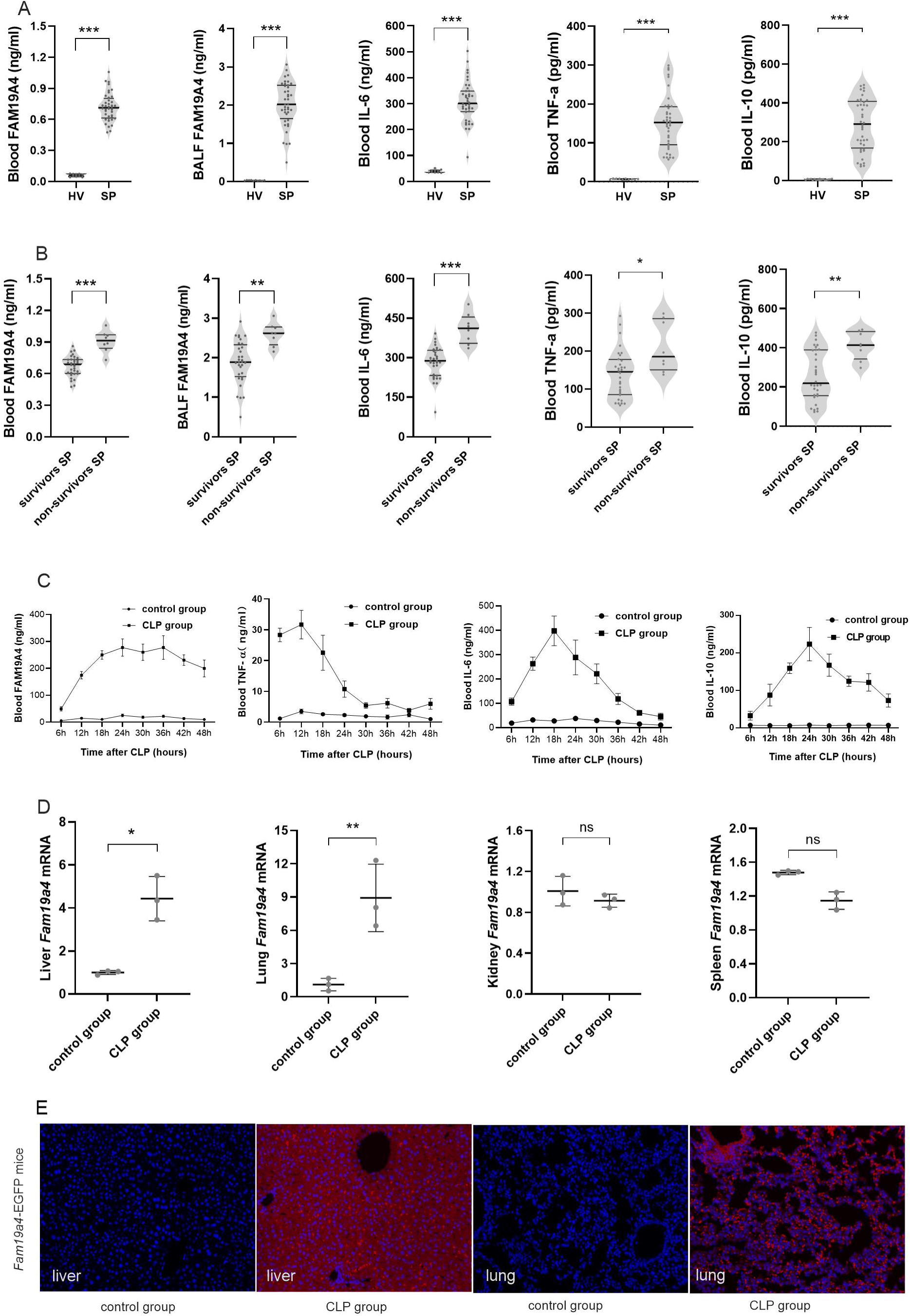
FAM19A4 expression is elevated in patients with sepsis and CLP-subjected mice. (**A–B**) FAM19A4 levels in serum and BALF samples from septic patients. Serum levels of IL-6, TNF-α, and IL-10 were used as reference markers of inflammatory response. (**A**) Patients with sepsis (SP, n = 40) vs. healthy volunteers (HV, n = 10); (**B**) survivor SP (n = 32) vs. non-survivor SP (n = 8). (**C**) FAM19A4, IL-6, TNF-α and IL-10 levels in mice serum 48 h after CLP. (**D**) *Fam19a4* expression in the liver, lung, kidney and spleen of mice 24 h after CLP. (**E**) *Fam19a4-*EGFP knock-in (KI) mice were used for CLP modeling. *Fam19a4* expression in the liver and lung was detected by in situ hybridization (200×). Blue dots, cell nuclei; red dots, *Fam19a4*. Results from one representative experiment out of three independent replicates are shown. (**A–D**) Data are presented as mean ± s.d. and analyzed by unpaired Student’s *t*-test. *P < 0.05, **P < 0.01, ***P < 0.001.

FAM19A4 is an evolutionarily conserved protein, and the amino-acid sequences of human FAM19A4 and its murine homologue are 93% identical [16]. Analysis of murine FAM19A4 mirrored our clinical findings. Mice with CLP-induced sepsis exhibited marked serum FAM19A4 elevation (Figure 1C). However, FAM19A4 expression in peritoneal lavage fluid showed no significant differences among the groups (Figure S2D). Tissue-specific analysis 24 h post-CLP demonstrated upregulated *Fam19a4* mRNA in liver/lung versus negligible baseline levels (Figure 1D). Analysis of *Fam19a4*-EGFP KI mice (Figure S1A, B) facilitated spatial resolution of *Fam19a4* expression patterns. In situ hybridization confirmed CLP-induced *Fam19a4* upregulation in hepatic/pulmonary tissues vs. minimal renal expression (Figure 1E, Figure S1C).

### 3.2 FAM19A4 deficiency reduces mortality and attenuates organ injury severity in CLP-subjected mice

FAM19A4 modulates innate immune cell function during inflammatory responses [18]. To investigate the role of FAM19A4 in sepsis, *Fam19a4*^−/−^ mice and WT littermate controls were subjected to CLP-induced sepsis. Complementary intervention involved intraperitoneal administration of recombinant FAM19A4 protein in WT mice at 1 and 3 h post-CLP to simulate hyperexpression. Genetic ablation of FAM19A4 significantly improved 7-day survival rates and attenuated weight loss compared to WT controls. Exogenous FAM19A4 supplementation did not alter survival outcomes or weight trajectories (Figure 2A, B).

**Figure 2.**
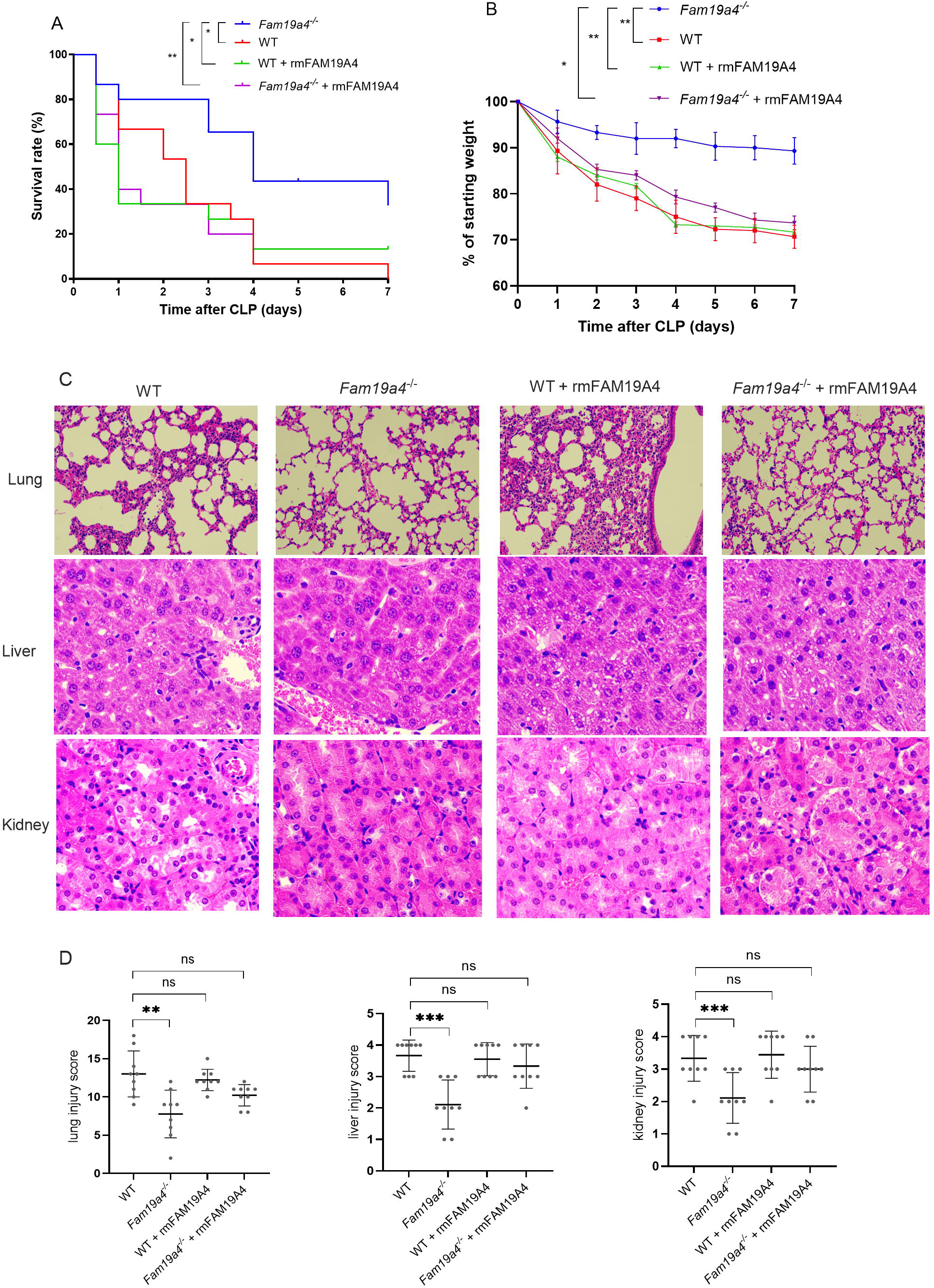
FAM19A4 deficiency reduces mortality and attenuates organ injury severity in CLP-subjected mice. (**A**) Survival rate and (**B**) body weight change (relative to initial weight) was monitored over time after CLP. (**C**) Representative H&E-stained section of lung, liver, and kidney tissues 24 h after CLP (200×). (**D**) Pathological injury scores of lungs, liver, and kidney for each group. Experimental groups: WT: WT mice subjected to CLP; *Fam19a4*^−/−^: *Fam19a4*^−/−^ mice subjected to CLP; WT + rmFM19A4: WT mice with CLP receiving intraperitoneal injection of rmFAM19A4 (100 µg/mouse) at 1 and 3 h after CLP; *Fam19a4*^−/−^ + rmFAM19A4: *Fam19a4*^−/−^ mice with receiving rmFAM19A4 as above; n = 3-6 per group. Data in (**B**) and (**D**) are presented as mean ± s.d. Image in (**C**) is representative of three independent experiments. *P < 0.05, **P < 0.01, ***P < 0.001; by log-rank test (**A**), two-way analysis of variance (ANOVA) (**B**), or Student’s *t*-test (**D**).

Liver, lung, and kidney tissue damage were assessed in mice at 24 h post-CLP by H&E staining and light microscopy (Figure 2C), with histopathological scoring of the damage (Figure 2D). CLP-subjected mice exhibited marked pathological alterations across multiple organs, including focal necrosis in liver lobules, increased hepatocyte destruction, and exacerbated renal tubular epithelial cell damage. In contrast, CLP-subjected *Fam19a4*^−/−^ mice displayed attenuated histological changes relative to those in WT mice. To validate these findings, recombinant FAM19A4 protein was administered intraperitoneally to *Fam19a4*^−/−^ mice at 1 and 3 h post-CLP, which restored severe pathological damage and elevated mortality rates. Transmission electron microscopy of liver and lung tissues 24 h post-CLP revealed vacuolar degeneration and rupture of intracellular mitochondria, endoplasmic reticulum, and other organelles in WT mice, whereas *Fam19a4*^−/−^ mice exhibited relatively mild ultrastructural lesions (Figure S2A). These findings suggest that FAM19A4 deficiency may mitigate sepsis severity and enhance clinical outcomes.

To investigate whether FAM19A4 directly modulates neutrophil function, thereby influencing recruitment of these cells to primary migration sites and subsequent control of microbial load, we analyzed changes in peripheral blood neutrophil counts and peritoneal bacterial burden in mice after manipulation of FAM19A4 expression. Following CLP, *Fam19a4*^−/−^ mice exhibited no significant increase in peritoneal bacterial colony counts relative to those in WT mice; however, intraperitoneal recombinant FAM19A4 (rmFAM19A4) protein administration significantly reduced peritoneal bacterial colonization in both WT and *Fam19a4*^−/−^ mice (Figure S2B). No significant differences in peripheral blood neutrophil counts were detected in any group (Figure S2C).

### 3.3 FAM19A4 deficiency decreases neutrophil and macrophage numbers in organs with reduced ROS and NET formation in CLP-subjected mice

Activation of the innate immune system during the early phase of sepsis is critical for bacterial clearance; however, the accumulation of inflammatory cells, such as neutrophils and macrophages, in non-infected distant organs, along with their activation and release of cytotoxic substances, significantly contributes to organ damage and dysfunction [26]. Liver and lung tissues were harvested from *Fam19a4*^−/−^ mice 24 h post-CLP. To quantify cell numbers, immunohistochemistry was performed using Ly6G^+^-labeled neutrophils and F4/80^+^-labeled macrophages. Neutrophil and macrophage numbers were significantly lower in liver and lung tissues from *Fam19a4*^−/−^ than in those of WT mice (Figure 3A, B).

**Figure 3.**
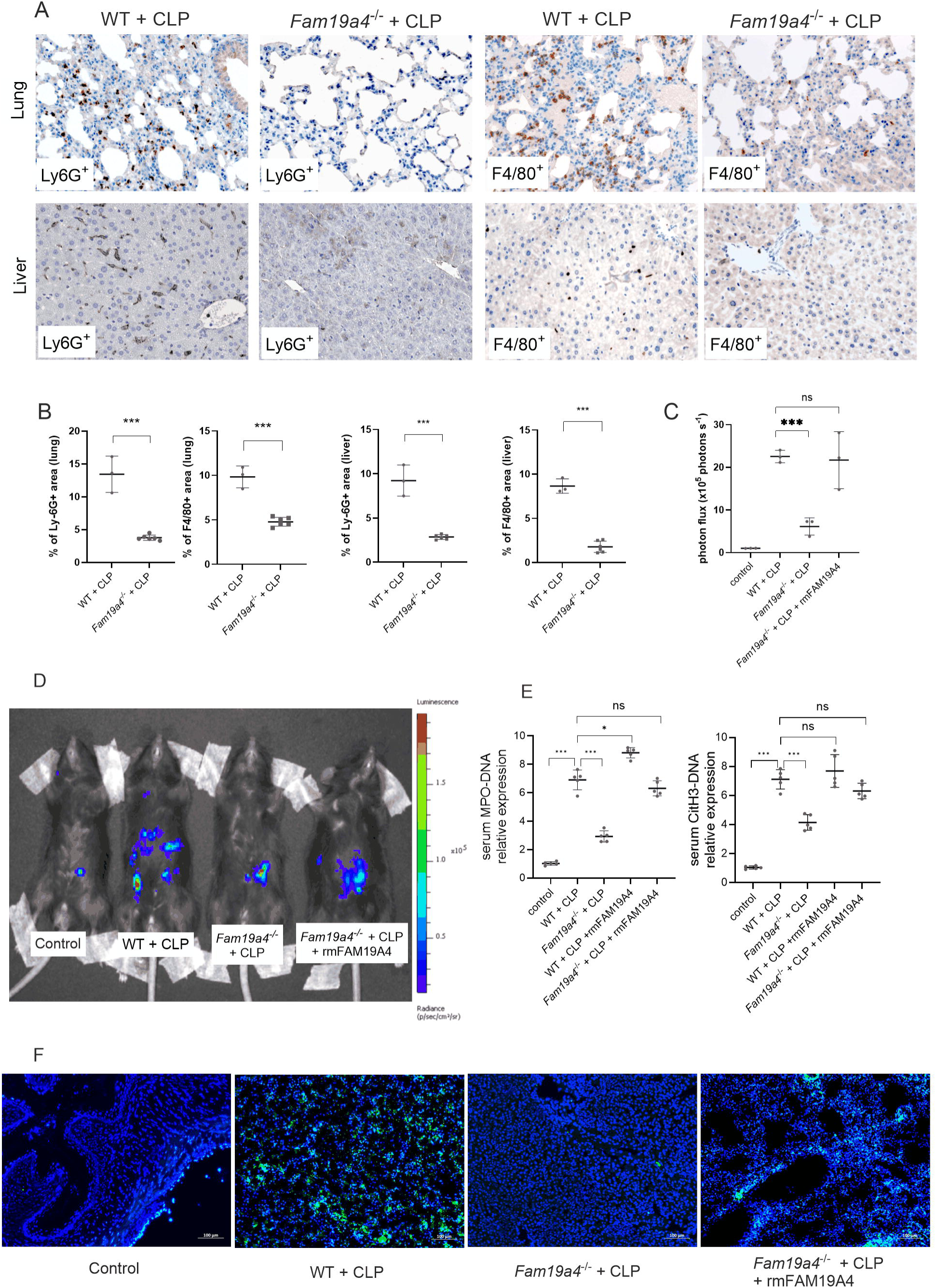
FAM19A4 deficiency decreases neutrophil and macrophage numbers in organs with reduced ROS and NET formation in CLP-subjected mice. (**A**) Representative immunohistochemical images of lung and liver sections stained for the neutrophil (Ly-6G) and macrophage (F4/80) in wild-type (WT) and *Fam19a4*^−/−^ mice 24 h after CLP (200×). (**B**) Quantification showing Ly6G^+^ and F4/80^+^ areas percentages in tissues (WT, n = 3; *Fam19a4*^−/−^, n = 6). (**C**) Average luminescence intensity in mice injected with L-012 24 h after CLP (n = 3 per group). (**D**) Representative bioluminescence images of ROS production. Pseudo colors represent photons/s/cm^2^/sr. (**E**) Circulating NET biomarkers (MPO-DNA and CitH3-DNA complexes) measured by ELISA. (**F**) Immunofluorescence imaging of NETs (CitH3, green) in lung tissue 24 h after CLP. Scale bars, 100 µm. Data in **B**, **C**, and **E** are presented as mean ± s.d.; *P < 0.05, **P < 0.01, ***P < 0.001; by Student’s *t*-test (**B**, **C**) or two-way ANOVA (**E**). Images in **A**, **D**, and **F** are from one representative experiment out of three independent replicates.

To investigate the effect of FAM19A4 on ROS levels in sepsis, L-012 (a luminol analog) was used to quantify systemic ROS production in CLP-subjected mice. At 24 h post-CLP, luminescence intensity was > 8-fold higher in WT mice than that in controls, with pronounced abdominal signal localization. This response was markedly attenuated in *Fam19a4*^−/−^ mice. Furthermore, intraperitoneal administration of rmFAM19A4 protein to WT mice at 1 and 3 h post-CLP (followed by L-012 labeling) resulted in enhanced ROS release compared to WT + CLP mice and restored peritoneal ROS levels in *Fam19a4*^−/−^ + CLP mice (Figure 3C, D).

ROS contribute critically to NET formation [27]. Dysregulated formation of NETs may cause cellular damage in adjacent tissues, thereby contributing to acute organ injury [12]. We next measured the extent of NET formation in both serum and organ tissue samples obtained from mice subjected to CLP, using CitH3-DNA and MPO-DNA complexes as key markers of circulating NETs. Exogenous administration of FAM19A4 by intraperitoneal injection led to a substantial elevation in serum levels of CitH3-DNA and MPO-DNA. In contrast, these biomarkers were significantly decreased in *Fam19a4*⁻*/*⁻ mice (Figure 3E). Next, we conducted immunofluorescence staining to assess the role of FAM19A4 in NET formation within the lungs.

Significantly fewer NETs (CitH3□ structures) were observed in *Fam19a4*□*/*□ mice than in WT mice at 24 h post-CLP (Figure 3F). Collectively, these data demonstrate that FAM19A4 deficiency attenuates NET formation in the serum and lung tissue of CLP-subjected mice.

### 3.4 FAM19A4 enhances neutrophil phagocytosis and ROS release in vitro

Neutrophils are the immune cells primarily responsible for ROS release during the early phase of sepsis [28]. To directly assess the functional impact of FAM19A4, LPS-pre-stimulated human neutrophils were treated with rmFAM19A4 in vitro, followed by evaluation of chemotaxis, phagocytosis, and ROS release. FAM19A4 exerted no significant chemotactic effect on neutrophils, regardless of LPS pre-stimulation (Figure S3). Confocal microscopy revealed that rmFAM19A4 enhanced neutrophil phagocytic capacity toward *E. coli*, with a more pronounced effect observed in LPS-pre-stimulated cells (Figure 4A-D). But blocking formyl peptide receptor 1 (FPR1) did not influence the effect of rmFAM19A4 in promoting phagocytic function (Figure S4).

**Figure 4.**
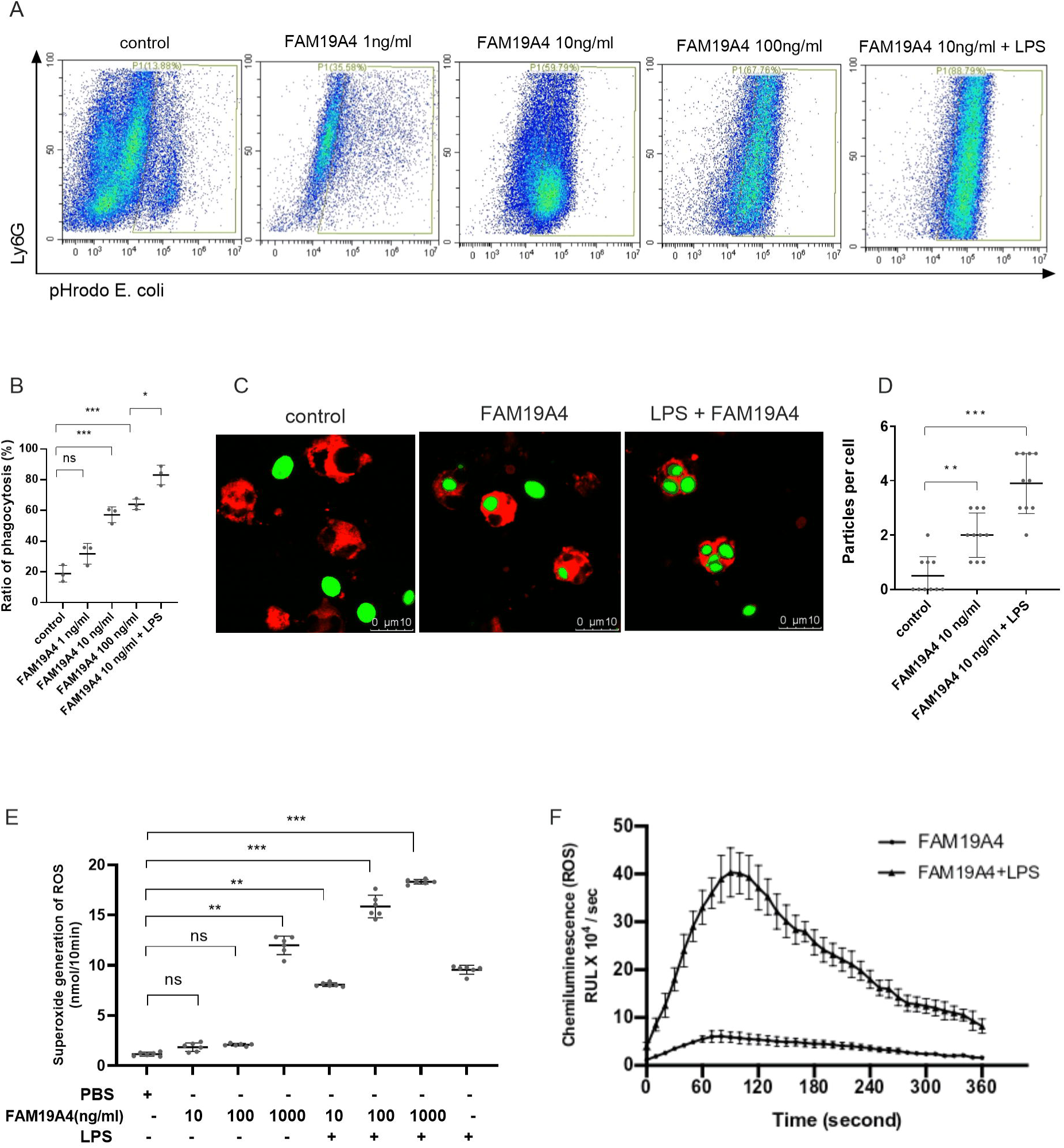
FAM19A4 enhances neutrophil phagocytosis and ROS release in vitro. (A, B) Neutrophils were treated with different concentrations of FAM19A4 protein, with or without LPS, and then incubated with pHrodo *E. coli* bioparticles. Phagocytosis was analyzed by flow cytometry. (A) Representative flow cytometry dot plots from three independent experiments. (B) Quantification of phagocytosis from pooled flow cytometry data. (C, D) Phagocytosis of pHrodo *E. coli* particles was observed by confocal microscopy. (C) Representative images. (D) Quantification of phagocytosed particles per neutrophil. (E) Superoxide release from neutrophils (1 × 10^6^ cells/mL), pre-incubated with PBS or LPS and then stimulated with FAM19A4 (1 to 100 ng/ml), was measured by ferricytochrome c reduction. (F) Total ROS production was detected using the luminol–peroxidase system in neutrophils treated with LPS or PBS, followed by DMSO or FAM19A4 (100 ng/mL). (B, D, E, F) Data are presented as mean ± s.d.; **P < 0.01, ***P < 0.001; by two-way ANOVA (B) or Student’s *t*-test (D, E, and F).

Neutrophils remain quiescent in peripheral blood, but bacterial components can prime their transition to an activation-ready state [29]. Differential ROS production across neutrophil activation states may contribute to dysregulated inflammatory responses in sepsis [30]. Neutrophils were treated with graded concentrations of FAM19A4, with experimental groups, including LPS- and PBS-pre-stimulated conditions. Extracellular superoxide anion (O^2·-^) generation was quantified via superoxide dismutase-inhibitable ferricytochrome C reductase assay, while total ROS levels were measured using luminol-enhanced chemiluminescence. In resting neutrophils (without LPS induction), low FAM19A4 concentrations exerted a minimal effect on ROS release, whereas higher concentrations significantly induced ROS production. Further, LPS-pre-stimulated neutrophils exhibited amplified ROS release, even at low FAM19A4 concentrations (Figure 4D, E).

### 3.5 p38 MAPK signaling is critical for FAM19A4-induced ROS in neutrophils

To elucidate the intracellular signaling pathways mediating FAM19A4 activity in human peripheral neutrophils, we performed bulk RNA-sequencing (RNA-seq) to identify transcription-level alterations in FAM19A4 target genes. The transcriptome profiles of PBS-treated controls and neutrophils exposed to graded concentrations of FAM19A4 for 2 h were compared. The differential gene expression profile, including both up-regulated and down-regulated transcripts in FAM19A4-treated neutrophils versus controls, is presented (Figure 5A). KEGG pathway analysis revealed marked enrichment of the MAPK signaling pathway following FAM19A4 exposure (Figure 5B), with heatmap visualization highlighting the 20 most significantly regulated transcription factors (Figure 5C). Pseudogenes and unclassified genes were systematically excluded from the analysis.

**Figure 5.**
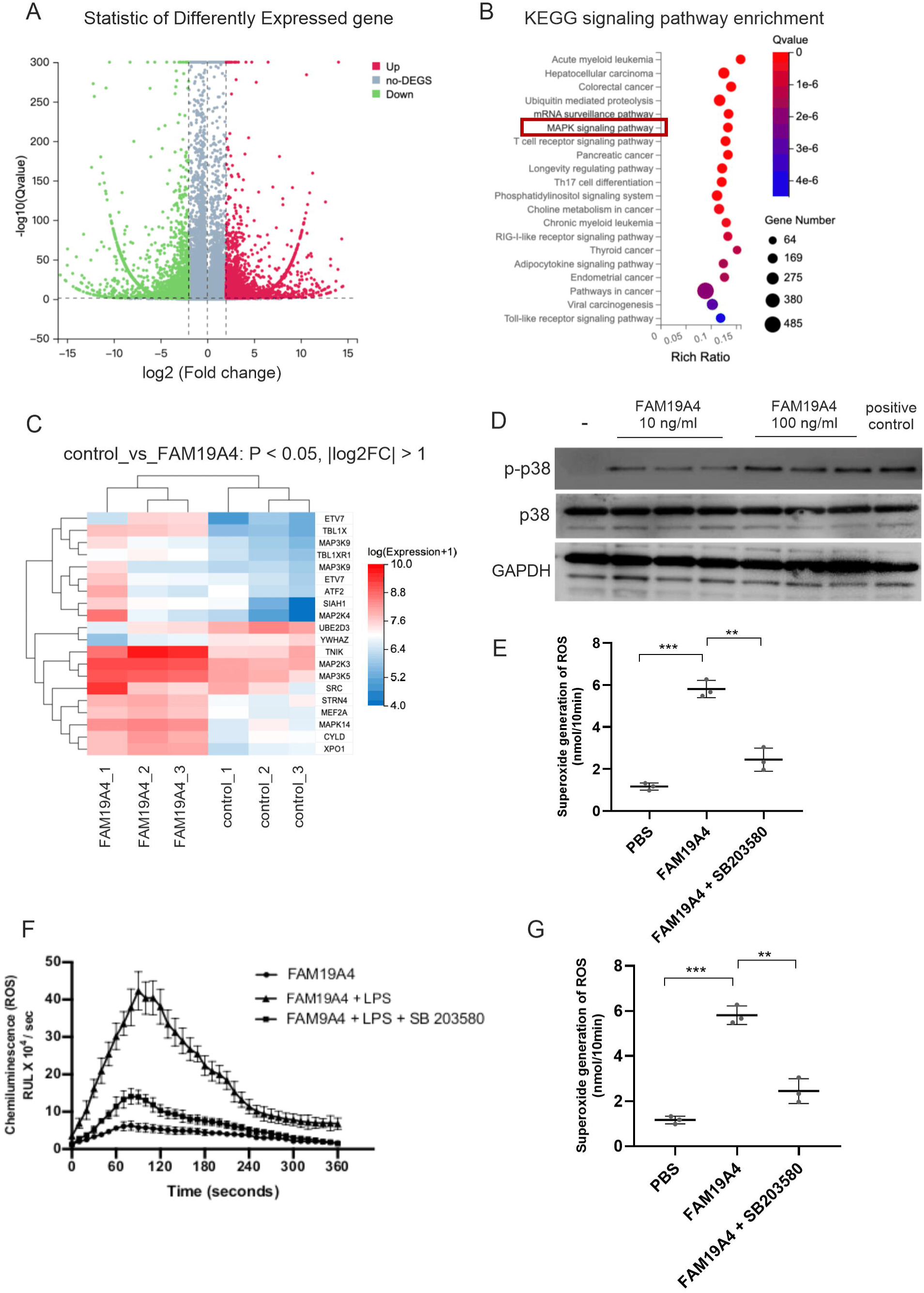
p38 MAPK signaling is critical for FAM19A4-induced ROS in neutrophils. (**A**) Volcano plot of differentially expressed genes in neutrophils treated with rmFAM19A4 (100 ng/ml, 2 h) versus PBS control. Red and green dots indicate up- and down-regulated genes, respectively. (**B**) KEGG enrichment analysis. Bubble size reflects gene number per pathway; color indicates Q value. (**C**) Heatmap of DEGs clustered by expression level (red: higher). (**D**) Western blot of p-p38 in isolated neutrophils stimulated with rmFAM19A4 (10 -100 ng/ml); GAPDH served as loading control. Image is representative of three experiments. (**E**) Superoxide release detected by ferricytochrome c reduction after pre-treatment with PBS or SB203580 (10 µM). (**F**) Total ROS measured via luminol–peroxidase system in neutrophils pre-treated with PBS, SB203580, or LPS. (**G**) Intracellular ROS levels (DCFH-DA MFI) in PBMCs from mice under four conditions: WT control, WT + CLP, Fam19a4−/− + CLP, and WT + CLP + SB203580. Data in (**E** - **G**) are mean ± s.d.; n = 3–5; **P < 0.01, ***P < 0.001, ****P < 0.0001 (Student’s *t*-test).

Guided by these findings and existing literature, we prioritized three MAPK sub-pathways (ERK1/2, p38 MAPK, and JNK) for functional validation. Using human peripheral neutrophils, we assessed signaling pathway activation post-FAM19A4 stimulation by western blot analysis. This approach identified significant enhancement of p38 phosphorylation following FAM19A4 treatment (Figure 5D, Figure S5). To establish functional causality, we compared

FAM19A4-treated neutrophils with those co-treated with the specific p38 inhibitor, SB 203580 [30]. Pharmacological inhibition of p38 substantially and significantly attenuated FAM19A4-induced ROS production relative to untreated controls (Figure 5E, F). We also intraperitoneally injected SB203580 to inhibit p38 MAPK function in the CLP model of WT mice, ROS release in the PBMCs of mice was significantly reduced (Figure 5G). Collectively, these data indicate that p38 MAPK signaling constitutes a key mechanistic pathway through which FAM19A4 potentiates neutrophil ROS release.

Moreover, clustering analysis further indicated an upregulation in Src kinase expression following FAM19A4 treatment (Figure 5C). According to a recent report^32^, ICAM-1 facilitates the release of ROS and NETs under septic conditions. Upon binding to LFA-1 (CD11a/CD18), ICAM-1 activates Src kinase and downstream signaling mediators, leading to p38 MAPK activation. To determine whether FAM19A4-enhanced phagocytosis is mediated through the ICAM-1 pathway, a cell-permeable penetrating-conjugated ICAM-1 C-terminal peptide (RQIKIWFQNRRMKWKKQRKIKKYRLQQAQ; Peptide 2.0) and a scrambled control peptide (RQIKIWFQNRRMKWKKVDDSDDFESVVSV; Peptide 2.0) were synthesized based on previously established methods [31]. These peptides were applied to whole blood samples to inhibit downstream ICAM-1 intracellular signaling. No significant difference in phagocytosis was observed between two groups (data not shown), suggesting that ICAM-1 is not involved in FAM19A4-mediated neutrophil phagocytosis.

## 4 Discussion

In this study, we identified that the secretory protein, FAM19A4, levels of which are elevated during the early innate immune response phase of sepsis, contributes to injury of remote organs, such as liver and lung, and is associated with poor prognosis. Under physiological conditions, FAM19A4 is negligibly expressed in most organs except the nervous system [16]. Recent studies have revealed roles in inflammatory responses [19, 32, 33]. Building on the findings of Wang et al. that FAM19A4 is a macrophage-derived secretory protein that induces ROS production [18], we hypothesized that it could play a critical role in acute infectious inflammation. Data from both clinical samples and CLP-induced mouse models confirmed that FAM19A4 is elevated in early sepsis, paralleling the surge of other cytokines such as IL-6, TNF-α, and IL-10. This temporal association suggests that FAM19A4 may contribute to the disruption of early immune homeostasis in sepsis.

Our study demonstrates that genetic ablation of FAM19A4 improves survival and ameliorates multi-organ damage in a CLP-induced sepsis model. Although FAM19A4 deficiency did not affect peripheral neutrophil counts or direct neutrophil chemotaxis in vitro, it significantly reduced neutrophil and macrophage counts in remote organs (liver and lung), suppressed systemic ROS production, and inhibited NET formation. Neutrophil-derived ROS represents one of the primary mechanisms for bacterial clearance. Accordingly, intraperitoneal administration of rmFAM19A4 in CLP mice - thereby elevating its local concentration - significantly reduced bacterial load in the peritoneal cavity. Notably, however, endogenous FAM19A4 expression was not markedly elevated in the peritoneal lavage fluid of CLP mice, and its genetic deletion did not compromise bacterial clearance at the primary infection site. These results suggest that FAM19A4 primarily exacerbates the detrimental aspects of neutrophil activity in sepsis - such as excessive ROS and NET-mediated tissue injury - rather than impairing their essential antibacterial role [10]. The spatial heterogeneity in FAM19A4 expression, together with its specific impact on ROS and NET generation in distant organs, highlights the therapeutic potential of targeting FAM19A4 to reduce tissue damage without weakening host defense.

The organ-specific neutrophil patterns likely arise from complex inflammatory networks involving both infiltrating and resident immune cells, and may also reflect contextual regulation of FAM19A4 signaling within different tissue microenvironments. Overall, FAM19A4 modulates neutrophil effector functions rather than recruitment. Future studies using single-cell sequencing could further clarify neutrophil subpopulations and their functional specialization in FAM19A4-mediated organ protection, particularly given emerging insights into resident neutrophil heterogeneity in sepsis pathophysiology.

FAM19A4 induces dose-dependent release of ROS from neutrophils: high concentrations directly trigger ROS production, while subthreshold levels of synergistically enhance LPS-primed ROS generation through amplified NADPH oxidase activation [6, 29, 34]. This priming phenomenon aligns with the presence of primed neutrophil subpopulations in septic patients [35]. Thus, sepsis-induced elevation of FAM19A4 in the blood and organs likely exacerbates oxidative stress in pre-activated neutrophils. We also observed increased neutrophil extracellular traps (NETs) formation, collectively promoting multiple organ dysfunction. ROS are key inducers of NETosis, in which PAD4 mediates chromatin decondensation via histone citrullination. Since most cytokines influence NET formation via PAD4, we hypothesize that FAM19A4 may also affect NETosis in a PAD4-dependent manner, warranting further mechanistic study.

The p38 MAPK pathway was identified as a key mediator of FAM19A4-induced ROS release, consistent with its established role in balancing antimicrobial responses and tissue injury in sepsis [36, 37]. However, given the complex network-based regulatory mechanisms inherent to sepsis, and in light of our results and previous literature, we suggest that the effects of FAM19A4 involve multifaceted network regulation beyond single-pathway modulation. Our bulk RNA-seq data revealed increased expression of Src kinase. Previous studies reported that FAM19A4 may promote macrophage chemotaxis and phagocytosis via FPR1[18], and that activated FPR1 can directly and atypically activate Src kinase, thereby promoting neutrophil phagocytosis and ROS release. However, our data showed that rmFAM19A4 enhanced neutrophil phagocytosis independently of FPR1 and had no effect on neutrophil chemotaxis. Consistent with this, Hoeffel et al. found that FAM19A4 exerted no chemotactic effects on mouse bone marrow-derived or peritoneal macrophages in vitro, suggesting that it regulates macrophage inflammatory profiles in an FPR1-independent manner [19].

Additionally, p38 MAPK acts as a downstream signaling component of ICAM-1, which activates Src kinase and subsequently triggers p38 MAPK activation. Nevertheless, prior studies have indicated that ICAM-1 intracellular signaling is not essential for Staphylococcus aureus phagocytosis or ROS generation in human neutrophils [31]. We also found that FAM19A4-enhanced phagocytosis was not dependent on ICAM-1. Further support that ICAM-1 signaling is not critical for these neutrophil functions. More in-depth studies are warranted to elucidate the molecular mechanisms by which FAM19A4 exerts its regulatory role in sepsis.

We observed that FAM19A4 selectively activates p38 MAPK without significantly affecting the JNK or ERK pathways. This selectivity is likely due to differences in upstream activation mechanisms, receptor preferences, and feedback regulation among MAPK family members. p38 MAPK is highly sensitive to stress and inflammatory stimuli (such as cytokines and ROS) and is primarily activated through the MKK3/MKK6 cascade [38, 39]. In contrast, JNK and ERK are more responsive to growth factors and differentiation signals, respectively [40, 41]. Structural features of p38, such as the TGY motif within its TxY activation loop, may facilitate its preferential phosphorylation by Src family kinases—consistent with our observation of upregulated Src expression [42]. Thus, the functional phenotype induced by FAM19A4 may reflect the assembly of specific receptor–signal complexes that precisely regulate the p38 pathway to coordinate ROS release and NET formation in neutrophils. The exact underlying mechanisms require further investigation.

In clinical practice, sepsis patients present at various stages due to differences in the interval between symptom onset and hospital admission. This study focused on early-phase sepsis, restricting enrollment to patients with a confirmed diagnoses within 7 days post-onset based on retrospective medical history. Consequently, an ideal dynamic blood sampling protocol (e.g., days 1, 3, 7, 14 and 28) was not feasible. The clinical cohort was relatively small (40 patients and 10 controls), and the clinical heterogeneity among sepsis patients may limit the generalizability of the relationship between FAM19A4 levels and clinical outcomes. Future studies should include larger population to validate these findings and further explore clinical correlations.

Our results demonstrate that the p38 MAPK inhibitor SB203580 can inhibit FAM19A4-induced ROS production, suggesting its potential therapeutic utility in sepsis. However, it has been reported that the effective dose of SB203580 is close to its toxic level in models of other diseases, indicating a narrow therapeutic window [43]. Moreover, the p38 MAPK pathway can be activated by multiple cytokines and membrane proteins in sepsis and represents a common node in the networked immune regulation of the condition [44]. The overall consequences of SB203580-induced inhibition within this complex regulatory network remain unclear. Therefore, further studies are warranted to evaluate the therapeutic potential of SB203580.

Our findings establish FAM19A4 as a key mediator in sepsis pathogenesis. Its effects appear closely integrated within the broader cytokine network, suggesting a complex role in the inflammatory cascade. Further studies—directly comparing FAM19A4 with other inflammatory mediators and utilizing multi-cytokine blockade models—are warranted to elucidate its unique versus synergistic functions and to evaluate its potential as a therapeutic target in sepsis.

## 5 Conclusions

This study reveals that elevated levels of the FAM19A4 cytokine exacerbate organ damage in sepsis, while its deficiency reduces mortality and tissue injury by diminishing neutrophil numbers in remote organs, thereby suppressing ROS formation. Mechanistically, FAM19A4 directly enhances neutrophil ROS production via p38 MAPK signaling without affecting chemotaxis or bacterial clearance. These findings identify FAM19A4 as a therapeutic target and advance understanding of dysregulated immune networks in sepsis.

## 7 Author Contributions

SL contributed to writing-original draft, validation and visualization. All of the authors contributed to review and editing. SL, FZ, HX and KZ contributed to data curation. SL and LJ contributed to formal analysis. SL FZ and TL contributed to funding acquisition. SL and LJ contributed to investigation. SL, FZ, LJ, HX, TL and WW contributed to methodology. SL contributed to project administration. SL, TL and TW contributed to resource. SL and TW contributed to supervision.

## 9 Acknowledgments

We thank Professor Wenling Han for her critical comments and assistance with the generation of KI mice. We further thank Yaxin Wang and Yunwei Lu for their assistance with the CLP methodology.

## 11 Data Availability Statement

The raw data supporting the conclusions of this article will be made available by the authors, without undue reservation.

