## Supplemental figures and table for "FAM19A4 Enhances Neutrophil Respiratory Burst via p38 MAPK in Lethal Sepsis"

### Supplementary Figures

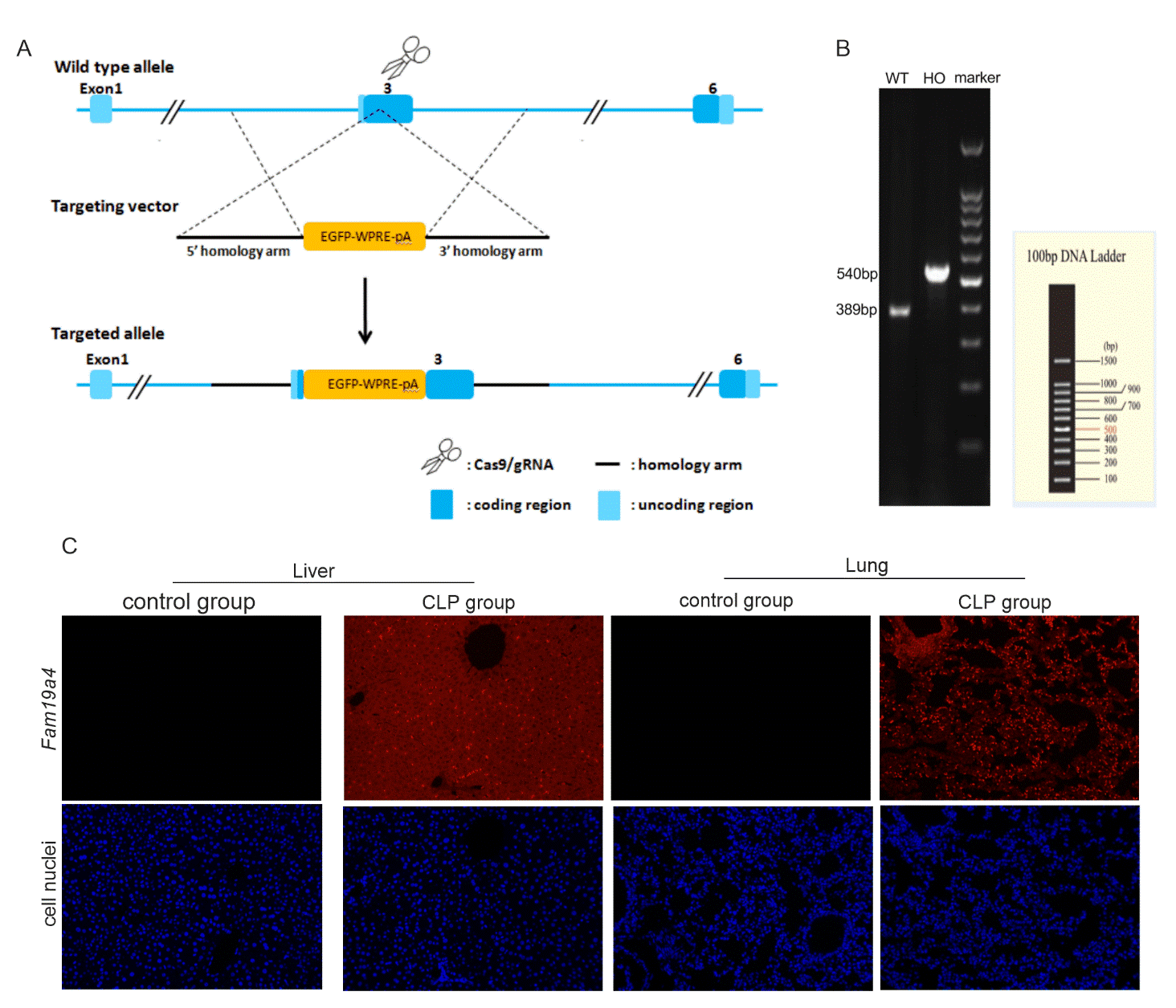

C

**Supplementary Figure S1.** FAM19A4 expression is elevated in patients with sepsis and CLP-subjected mice. (**A**, **B**) Generation and characterization of *Fam19a4*-EGFP mice. (**A**) Schematic representation of *Fam19a4*-EGFP BAC-based strategy used to trigger homologous recombination in the *Fam19a4* locus to generate *Fam19a4* knock-in mice. (**B**) Gel image showing genotype analysis of F1 heterozygous mice. Marker: 100bp Plus DNA Ladder (Transgen; BM311-01). (**C**) Representative in situ fluorescence imaging of Fam19a4 expression (red) and nuclei (blue) in liver and lung sections (200×). Results are from one of three independent experiments.

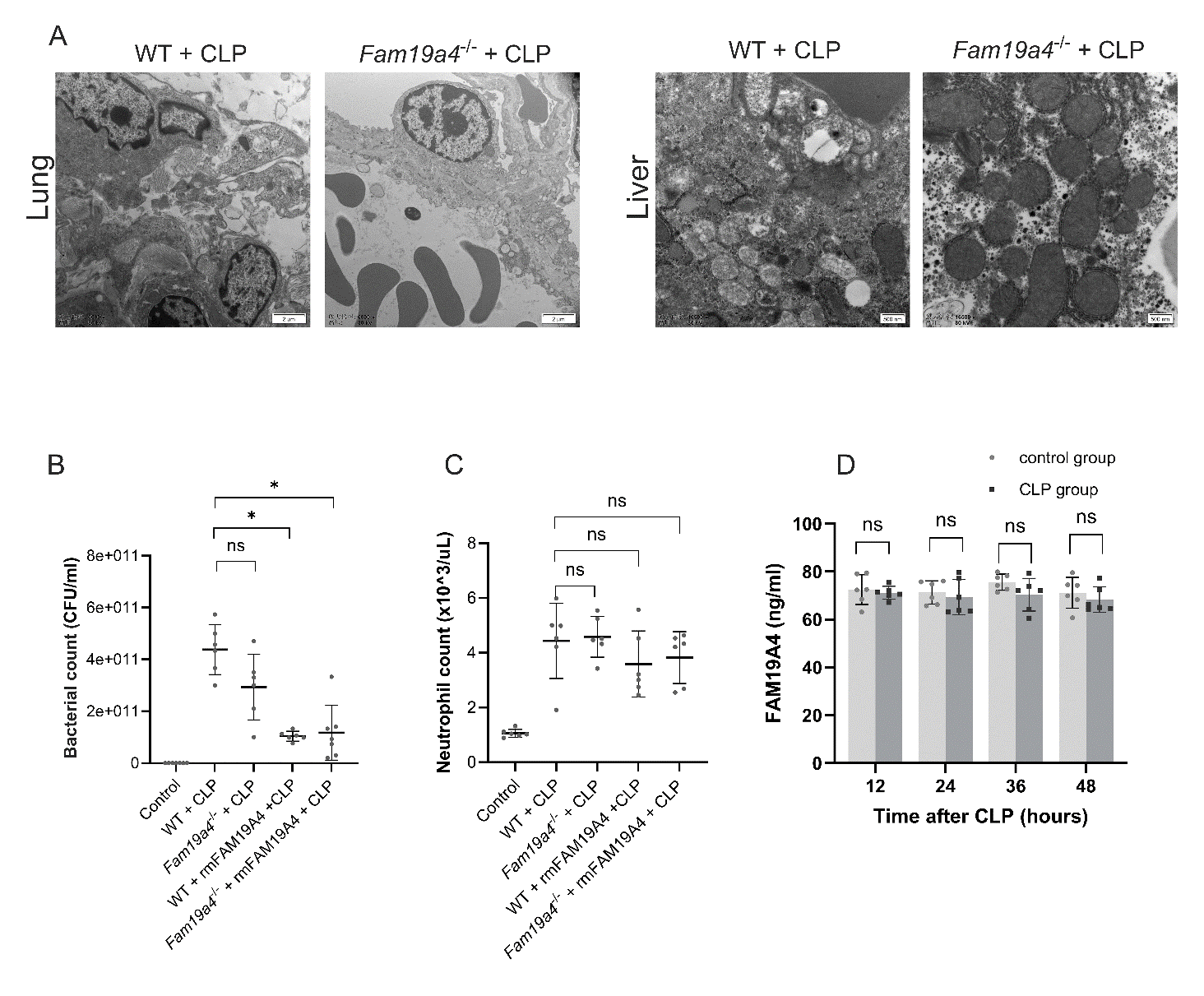

**Supplementary Figure S2.** Absence of FAM19A4 reduces organ damage but does not affect bacterial clearance or blood neutrophil counts in CLP-induced septic mice. (**A**) Representative transmission electron microscopy images of liver and lung tissues from WT and *Fam19a4*-/- mice 24 h after CLP. Scale bars, 2 μm (lung), 500 nm (liver). Image is from one of three independent experiments. (**B**) Bacterial load in peritoneal lavage fluid 24 h after CLP, determined by serial dilution and CFU counting on nutrient agar plates. (**C**) Peripheral blood neutrophil counts measured using a hematology analyzer (Pukang PE-6100). Control mice underwent sham laparotomy; treated mice received rmFAM19A4 (100 μg/mouse) intraperitoneally at 1 h and 3 h post-CLP. n = 3–6 per group. (D) FAM19A4 protein expression in peritoneal lavage fluid. Data are presented as mean ± s.d. *P < 0.05, **P < 0.01, ***P < 0.001 (Student’s *t*-test).

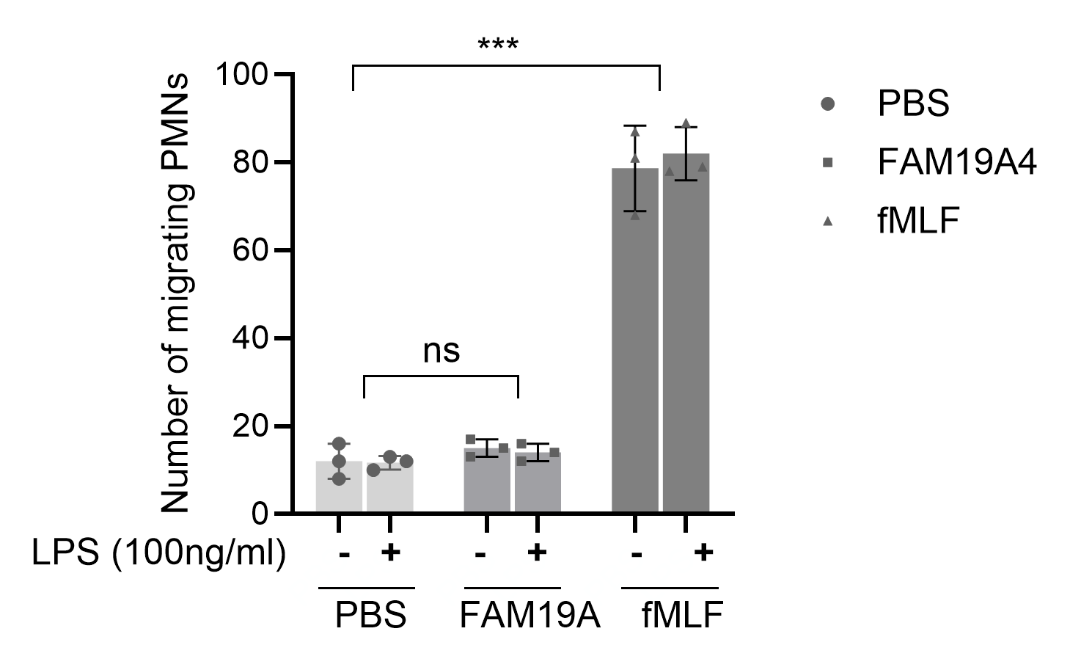

**Supplementary Figure S3.** FAM19A4 did not affect neutrophil chemotaxis, as assessed using a Transwell chamber assay. Human neutrophils were pretreated with PBS or LPS (10 μg/ml, 30 min). N-Formylmethionine-leucyl-phenylalanine (fMLF) was used as a positive control. Data are presented as mean ± s.d. from three independent experiments. ***P < 0.001 vs. PBS group (Student’s *t*-test).

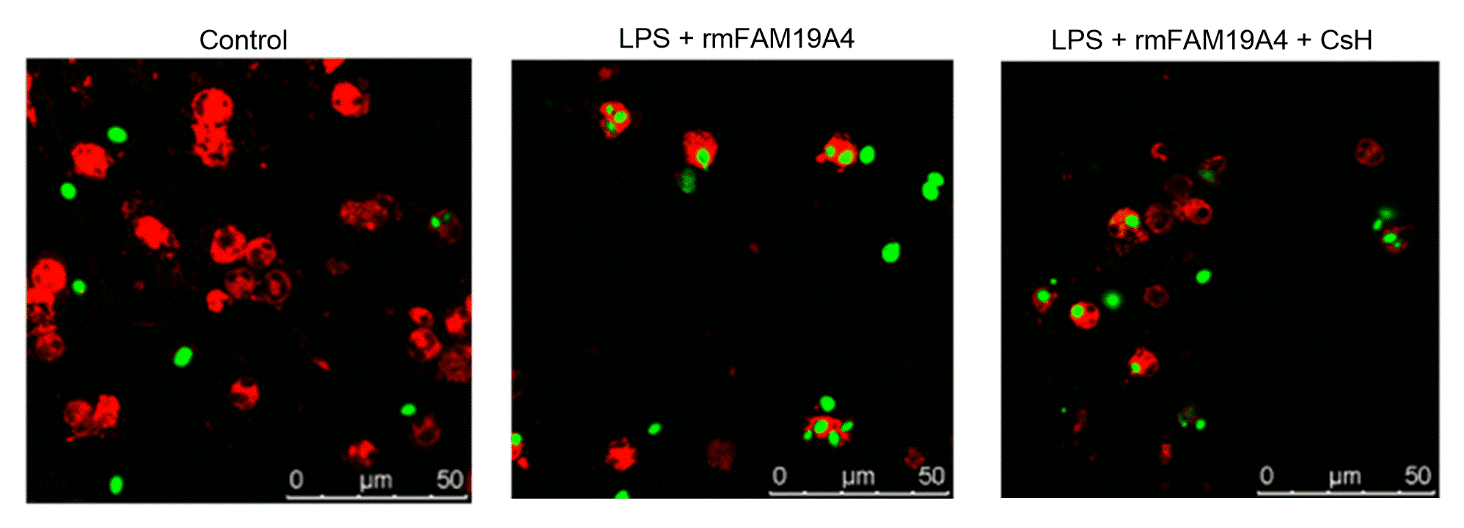

**Supplementary Figure S4.** Cyclosporine H (CsH) does not block rmFAM19A4-mediated enhancement of neutrophil phagocytic function. Numbers of FITC-*E. coli* particles phagocytosed by individual neutrophils were observed via confocal microscopy. Scale bars, 50 μm. CsH was used as an FPR1 blocker. Representative images from one of three independent experiments are shown.

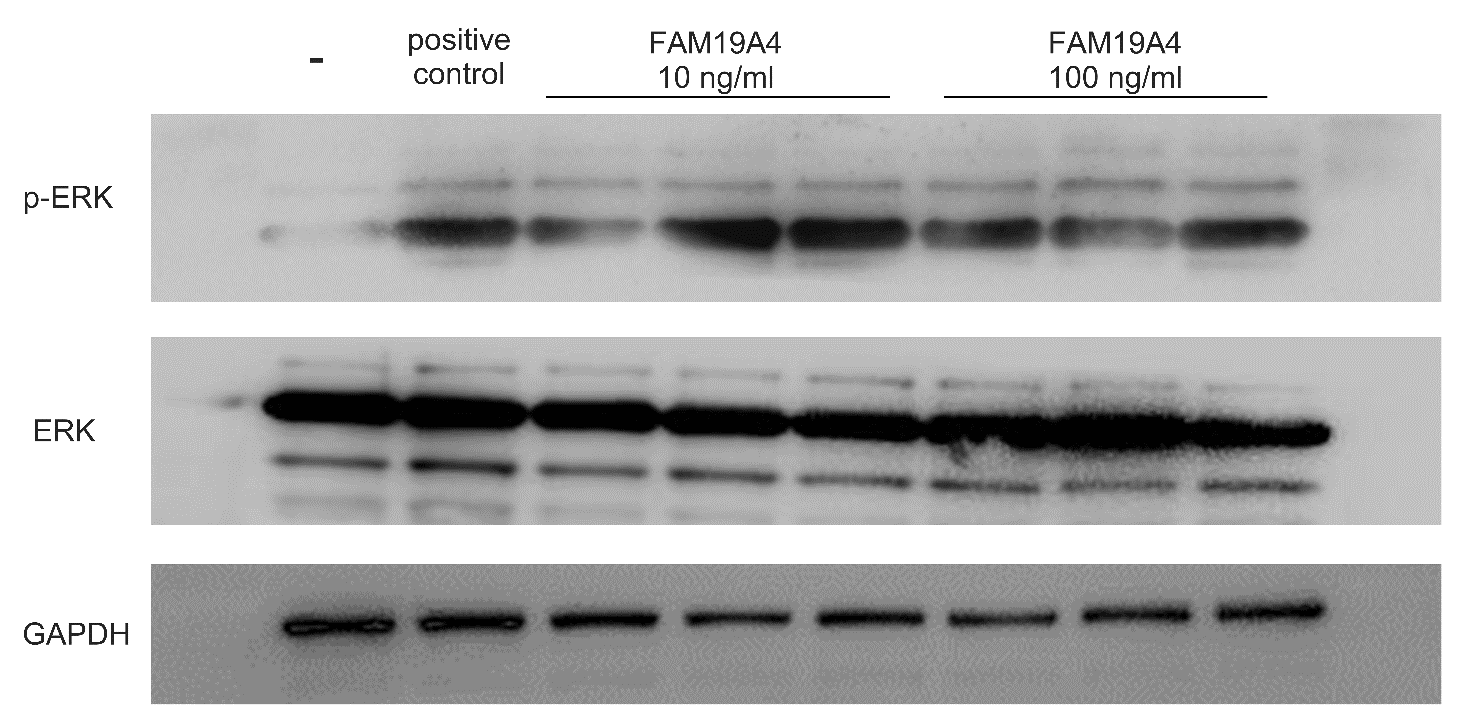

**Supplementary Figure S5.** Representative western blot of phosphorylated ERK in neutrophils treated with rmFAM19A4 protein (10–100 ng/ml); GAPDH was used as a loading control. Results of one representative experiment from three independent experiments are shown.

### Supplementary Table1. Log2 fold changes and adjusted Q-values of MAPK signaling kinases

| Gene | log2Ratio | Q value |
| --- | --- | --- |
| p38a | 0.535801434 | < 0.0001 |
| ERK1 | -0.016395313 | 0.567609706 |
| ERK2 | 0.017219397 | 0.503428582 |
| JNK1 | 0.011325613 | 0.954990391 |
| JNK2 | 0.171596987 | 0.195967653 |
| JNK3 | 0.225826128 | 0.294828663 |
